# A novel genetic strategy to interrogate an unknown phenotypic modifier: an Sdhc KO-Robertsonian mouse with a semi-homologous chromosome develops papillary thyroid carcinoma-like tumours

**DOI:** 10.1101/2025.10.31.685748

**Authors:** Jean-Pierre Bayley, Heggert Rebel, Peter Devilee

## Abstract

*SDHD* and *SDHAF2* pathogenic variants confer a remarkable parent-of-origin tumour risk for the neuroendocrine tumours paraganglioma and pheochromocytoma. Paternally transmitted variants cause tumours but maternally transmitted variants do not. The Hensen hypothesis asserts that loss of an (unknown) imprinted gene(s), together the remaining wildtype SDH gene, is a prerequisite for tumour formation. This study had three objectives, first, as a test of the Hensen model, second, as a potential paraganglioma model, and finally, as a test of chromosomal configuration to interrogate large genomic regions carrying an unknown phenotypic modifier. We crossed an SDH gene knockout line to a Robertsonian (Rb) chromosome line harbouring the gene imprinting centre implicated in human tumourigenesis, to create a metacentric chromosome with characteristics of human chromosome 11. Distinct phenotypes were noted in various cohorts. In heterozygote Rb mice we noted both weight gain and frequent immune activation. In Sdhc knockout mice with both heterozygous and homozygous Rb chromosomes, thyroid abnormalities, including papillary thyroid carcinoma-like tumours, were common due to apparent synergy between the Sdhc KO and the Rb chromosome. We also found a single case of bilateral pheochromocytoma in which loss of Sdhc was not the driver. Although few studies of Robertsonian chromosomes in the mouse have addressed pathology or phenotype, this study suggests that chromosomal structure can dramatically impact clinical phenotype.

## Introduction

Paragangliomas and pheochromocytomas/extra-adrenal paragangliomas (PPGL) are closely-related neuroendocrine tumours associated with the parasympathetic and sympathetic nervous systems, respectively. Parasympathetic paragangliomas are most commonly found in the head and neck region, where the most frequent site is the carotid body at the bifurcation of the internal and external carotid arteries. Pheochromocytomas arise in the adrenal medulla and sympathetic paragangliomas in the extra-adrenal sympathetic paraganglia [1]. Most cases of hereditary paraganglioma are attributable to genes encoding subunits of mitochondrial tricarboxylic acid cycle enzyme succinate dehydrogenase (SDH), including *SDHA, SDHB, SDHC, SDHD*, and the assembly factor gene *SDHAF2*. The SDH protein complex consists of two catalytic subunits, SDHA and SDHB, and two membrane-spanning subunits, SDHC and SDHD, each encoded by a specific gene. *SDHB* and *SDHC* genes are both found, widely separated, on human chromosome 1. *SDHA* is located on human chromosome 5, while *SDHD* is found on human chromosome 11.

Unlike pathogenic variants in other SDH subunit genes, variants in *SDHD* and *SDHAF2* (also located on chromosome 11) show a remarkable parent-of-origin dependent tumourigenesis in which tumour formation occurs almost exclusively following paternal transmission of the variant. A large majority of carriers of a paternally transmitted variant will develop a tumour at some stage during their lifetime, whereas virtually all maternal carriers will remain tumour-free for life. The absence of tumourigenesis in carriers of maternally transmitted *SDHD* variants initially suggested a maternally imprinted gene as the underlying cause of the tumour [2-4].

However, following identification of *SDHD* in 2000 it was quickly shown that the gene displays biallelic expression in a range of foetal and adult tissues, refuting the idea of direct imprinting of *SDHD* [5, 6]. This and the uniform loss of the entire maternal copy of chromosome 11 in human tumours carrying variants in *SDHD* or *SDHAF2* led to the formulation of the ‘Henson model’ [7]. This hypothesis proposes that a prerequisite for development of *SDHD- and SDHAF2*-related paraganglioma, besides a variant in the respective gene, is loss of a paternally-imprinted, maternally-expressed modifier gene likely found in the major gene imprinting centre on human chromosome 11p. Loss of chromosome 11, and especially 11p, is also a characteristic of *SDHB*-related tumours [8]. It is important to recall that the SDHD and SDHAF2 proteins play disparate roles in SDH complex function. SDHD is a transmembrane subunit, whereas SDHAF2 is an assembly factor that interacts transiently with SDHA during SDH complex assembly. This further supports the idea that gene location rather than protein function is likely the determining factor in the common inheritance pattern. The basis of the Hensen model is that chromosomal location, rather than the specific SDH gene, determines the characteristic pattern of tumour risk.

Extending the Henson hypothesis to the mouse, we predict that a mouse will develop paraganglioma only when SDH gene function is lost together with the modifier gene located in the equivalent imprinting centre found on mouse chromosome 7. We previously tested one candidate modifier gene (H19) in a cross to an Sdhd KO but that mouse did not develop PPGL [9]. The mouse imprinted region on chromosome 7 contains approximately the same genes as those found in the human imprinting centre. Imprinted genes differ from other genes in that only one parental copy is expressed or predominantly expressed, and most imprinted genes show similar patterns of imprinting in mouse and man [10]. To develop a ‘Hensen model’ in the mouse we crossed an Sdhc knockout mouse line with a mouse line carrying a Robertsonian (Rb) chromosome [Rb(1:7)]. Rb chromosomes are stable translocations of normally telocentric mouse chromosomes and thus resemble human metacentric chromosomes [11]. Diverse and multiple translocations are found in wild populations of subgenus Mus (incl. *Mus musculus domesticus*), some of which have been crossed with laboratory mouse strains and are commercially available. An Rb stain combining Sdhd (mouse Ch.9) and the mouse imprinting centre on mouse chromosome 7 was not available, so we instead selected the Rb(1:7) strain which harbours a single Robertsonian chromosome that combines Sdhc (Ch.1) with the mouse imprinting center (Ch.7). As outlined above, we predict that combining functional loss of any SDH gene with loss of the imprinted region may lead to tumour initiation.

The current lack of mouse models for SDH-related paraganglioma is a major obstacle to research. Numerous mouse models have been described over the past two decades [12], but none have developed paraganglioma to date. In view of the Hensen model, an unknown gene (or genes) playing an essential permissive role in human PPGL may be the missing piece in the puzzle of why mice fail to develop SDH-related paraganglioma. This approach serves a triple function as evidence supporting or refuting the Hensen model, as a potential paraganglioma model, and as a test of the use of chromosomal configuration to interrogate large genomic regions carrying an unknown phenotypic modifier.

## Methods

### Cohort generation

The C57BL/6N-SdhcTm1a(EUCOMM) Wtsi mouse, obtained from the Sanger Center, UK, on behalf of the European Mutant Mouse Archive, was developed by the European Conditional Mouse Mutagenesis (EUCOMM) Program and has been extensively phenotyped by the International Mouse Phenotyping Consortium (IMPC), which reported homozygous embryonic lethality, as well as increased body weight and thrombocytosis in heterozygotes. These animals carry a LacZ cassette inserted into intron 3 of Sdhc that causes skipping of exon 4. Skipping in the provided animals was confirmed by RT-PCR, and cross breeding ratios of heterozygote mice indicated homozygote lethality. The Rb mouse line, B6Ei.Cg-Rb(1.7)1Rma/J [Universita di Roma Rb(1:7)] was obtained from the Jackson Laboratory. This Rb strain carries a single translocation of chromosome 1 and chromosome 7.

Male mice carrying a heterozygous Rb chromosome [Rb/wt] were initially bred to wild type C57BL/6N females, and mice heterozygote for the Rb chromosome were then interbred to obtain heterozygote [wt/Rb] and homozygote mice [Rb/Rb]. Mouse cohorts were generated by initial crossing to ingress the Sdhc KO allele to the Rb background: Sdhc+/-x Rb1:7(Sdhc+/+)/Rb Rb1:7(Sdhc+/+), resulting in Sdhc-/RbSdhc+ mice. Sdhc-/Rb1:7(Sdhc+) mice were then crossed with Rb1:7(Sdhc+)/Rb Rb1:7(Sdhc+) mice and bred as available. Taking into account known transmission distortion in Rb mice (around 10%) and high recombination rates due to chromosomal distance, we expected an efficiency of ∼10% to generate Rb1:7(Sdhc-)/Rb1:7(Sdhc+) mice. Subsequent crosses of Rb1:7(Sdhc-)/Rb1:7(Sdhc+) x Rb1:7(Sdhc-)/Rb1:7(Sdhc+) generated further experimental and control cohorts. Mice were genotyped by PCR for the presence of the mutant Sdhc allele and results were compared to metaphase spreads (using standard protocols) from the same animals to determine combined genetic and chromosomal status.

### Animals and pathology

Mice were managed according to guidelines of the the Animal Experiment Committee of Leiden University Medical Center (DEC nr.12069), and housed under standard conditions (temperature of 22 ± 1.5 °C, 12-h light/12-h dark cycle) with constant access to food and water.

Mice showing serious signs of distress were sacrificed and investigated for signs of general disease in all organs, with a special focus on paraganglioma-related areas including the carotid bifurcation and the adrenal gland. Mice were followed biweekly and the experiment was terminated at approximately two years. Mouse pathology was assessed by the investigators, by an experienced human pathologist and by an experienced LUMC animal pathologist. Tumours and selected pathological tissues were snap frozen and preserved as formalin-fixed paraffin-embedded (FFPE) tissue.

### Statistics

Standard descriptive statistics were used to analyse data, including Student’s T test and 2x2 contingency tables, where appropriate (Graphpad, USA). All outcomes were corrected for multiple testing.

## Results

### Experimental cohorts

In total we developed 7 cohorts to test the Hensen model (Figures 1a & 1b), consisting of 3 control cohorts: (1) Sdhc +/-on the BL6 background (no Rb), (2) an Sdhc +/+ line with a single Rb chromosome, and (3) an Sdhc +/+ line with two Rb1:7 chromosomes. The experimental cohorts (4-7) all included Sdhc+/-with either one or two Rb chromosomes (to address possible differences in chromosomal stability). These crosses were conducted via either the female (4&6) or male (5&7) line in order to model human patterns of inheritance. The two cohorts carrying the paternally-transmitted Sdhc KO (4&6) simulate patients at risk of tumour development due to a paternally-inherited *SDHD* or *SDHAF2* variant (Figure 1c). This genetic configuration uniting Sdhc (Ch.1) and the imprinting centre (Ch.7) loosely models human chromosome 11, with the caveat that other intervening chromosomal regions differ from human chromosome 11 (Figure 2). As tumour development in this Sdhc-Rb model is dependent on the spontaneous loss of the normal copy of the chromosome, we generated and followed a large experimental group of 126 mice, including 80 in the paternally-transmitted cohorts. As clinical detection of PPGL in man typically occurs around the ages of 35-40 years, we followed these mice until aged (mean 70 weeks, range 17-152 weeks).

**Figure 1.**
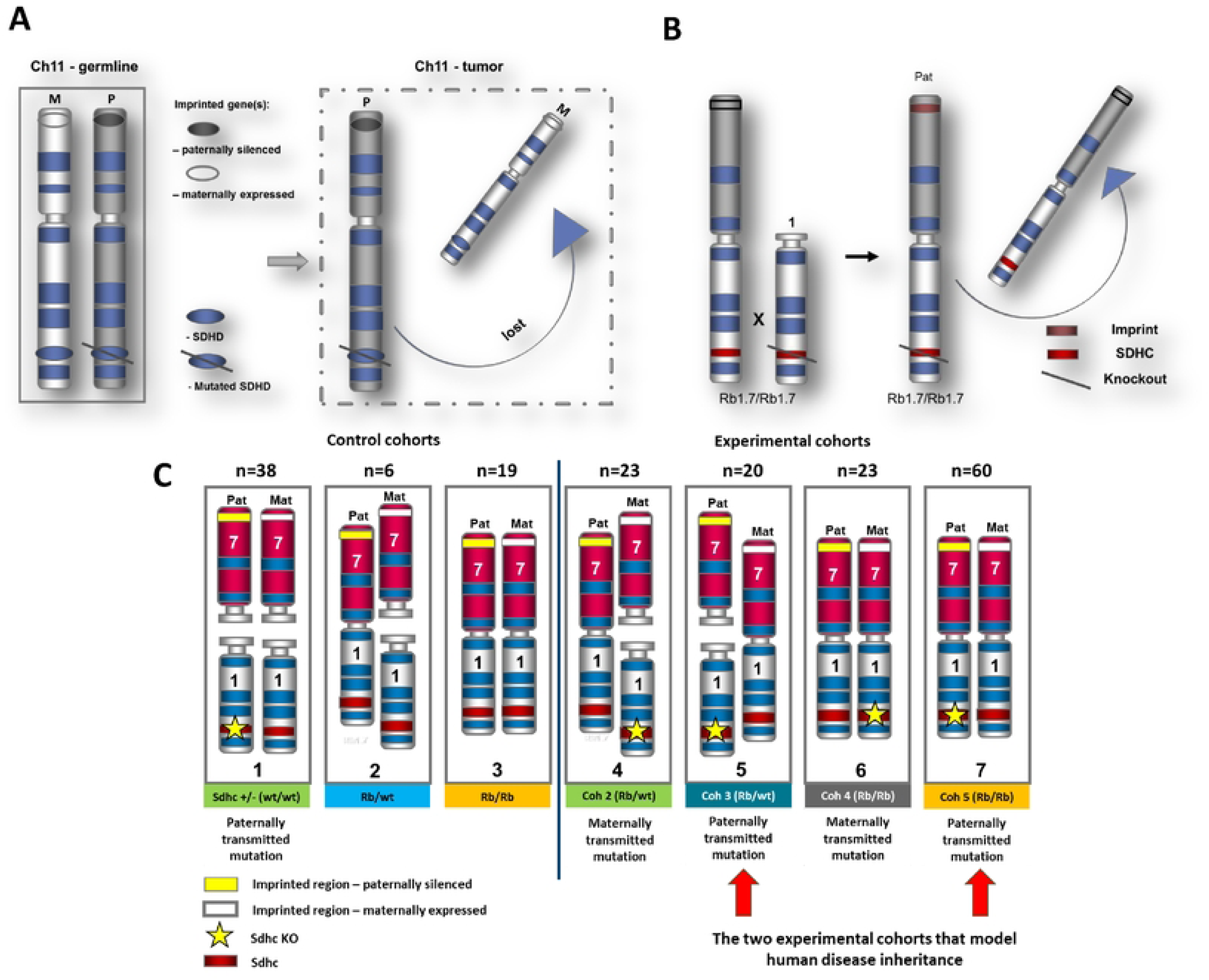
(A) The ‘Hensen model’ in outline. Loss of an imprinted gene in the tumour, together with the normal gene copy of SOHO, leads to tumour formation. (m - maternal, p - paternal) **(B).** The ‘Hensen model’ in the mouse. The Rbl:7 Robertsonian chromosome carries a PPGL tumour suppressor gene (Sdhc - chromosome 1 (red) one copy mutated) and the mouse equivalent of the human imprinting centre on human chromosome 11 (open/closed box on (grey) mouse chromosome 7). The depicted scenario was used to generate cohorts 4 and 5, which carried two copies of the Rbl:7 chromosome. **(C)** Strategy to create an (approximate) metacentric homolog of human chromosome 11 in the mouse. Sdhc +/-Rbl:7 experimental cohorts in outline. These cohorts were designed to model the parent-of-origin inheritance effect in the mouse. Cohorts 1, 4-7 all carry a heterozygous knockout (null allele) of Sdhc. According to this model, both functional gene copies (Sdhc + unknown imprinted modifier) should be lost following an LOH event, leading to tumorigenesis. Cohort 7 most closely resembles the Hensen model in man. Cohorts 4 and 5 were created assess possible differences in chromosomal instability due to configuration of Robertsonian chromosomes. The cohorts with maternally-inherited mutations acted as controls for the Hensen model. (Mat=maternal, Pat=paternal).

**Figure 2.**
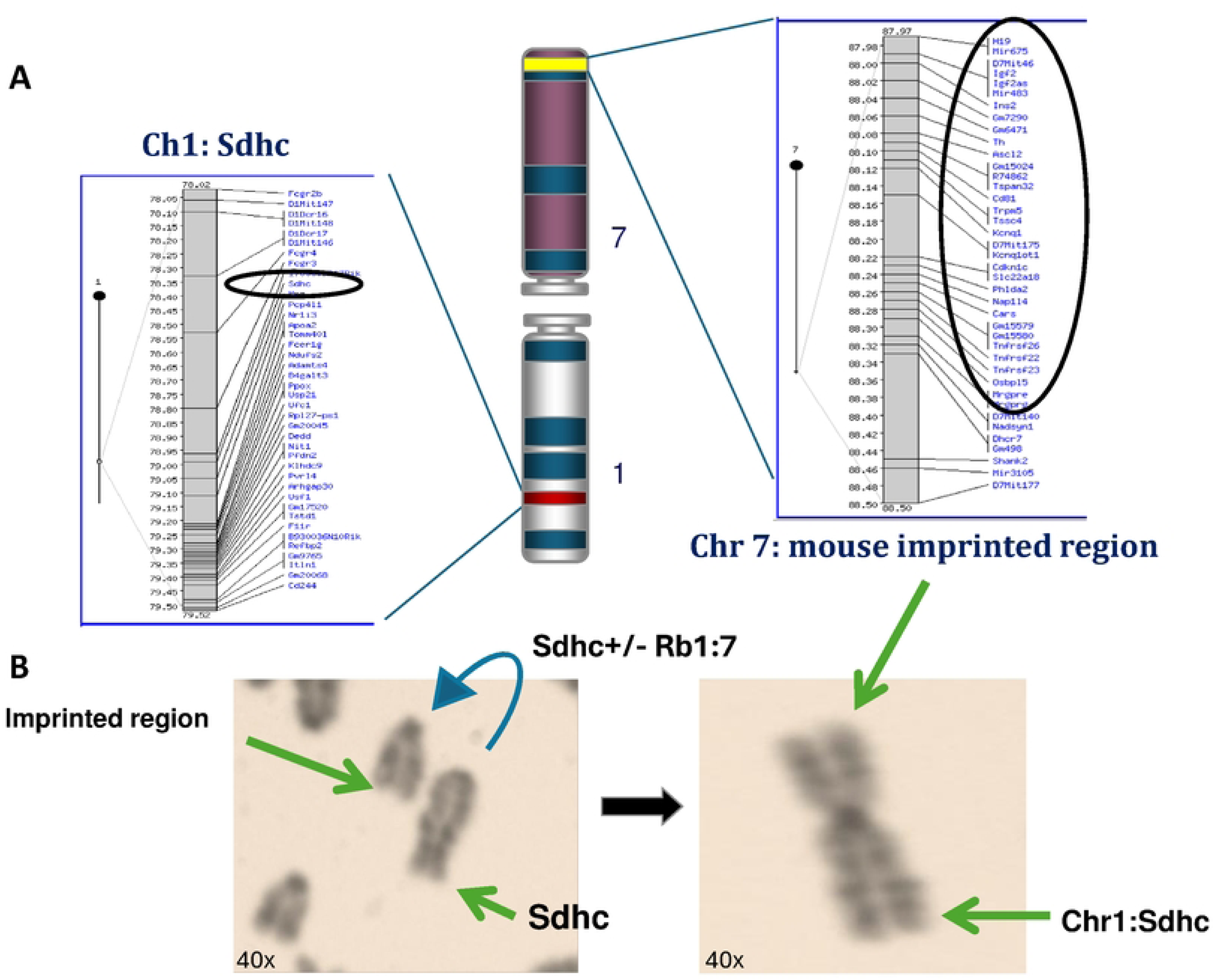
(A) Human chromosome 11 and mouse chromosome 7 share twenty genes that are reportedly imprinted in one or both species (https://www.geneimprint.com/site/genes-by-species). See also supplementary data 1 **(B).** A metaphase spread from a B6Ei.Cg-Rb(l.7)1Rma/J mouse illustrating the visual appearance of a Robertsonian chromosome.

**Figure 3.**
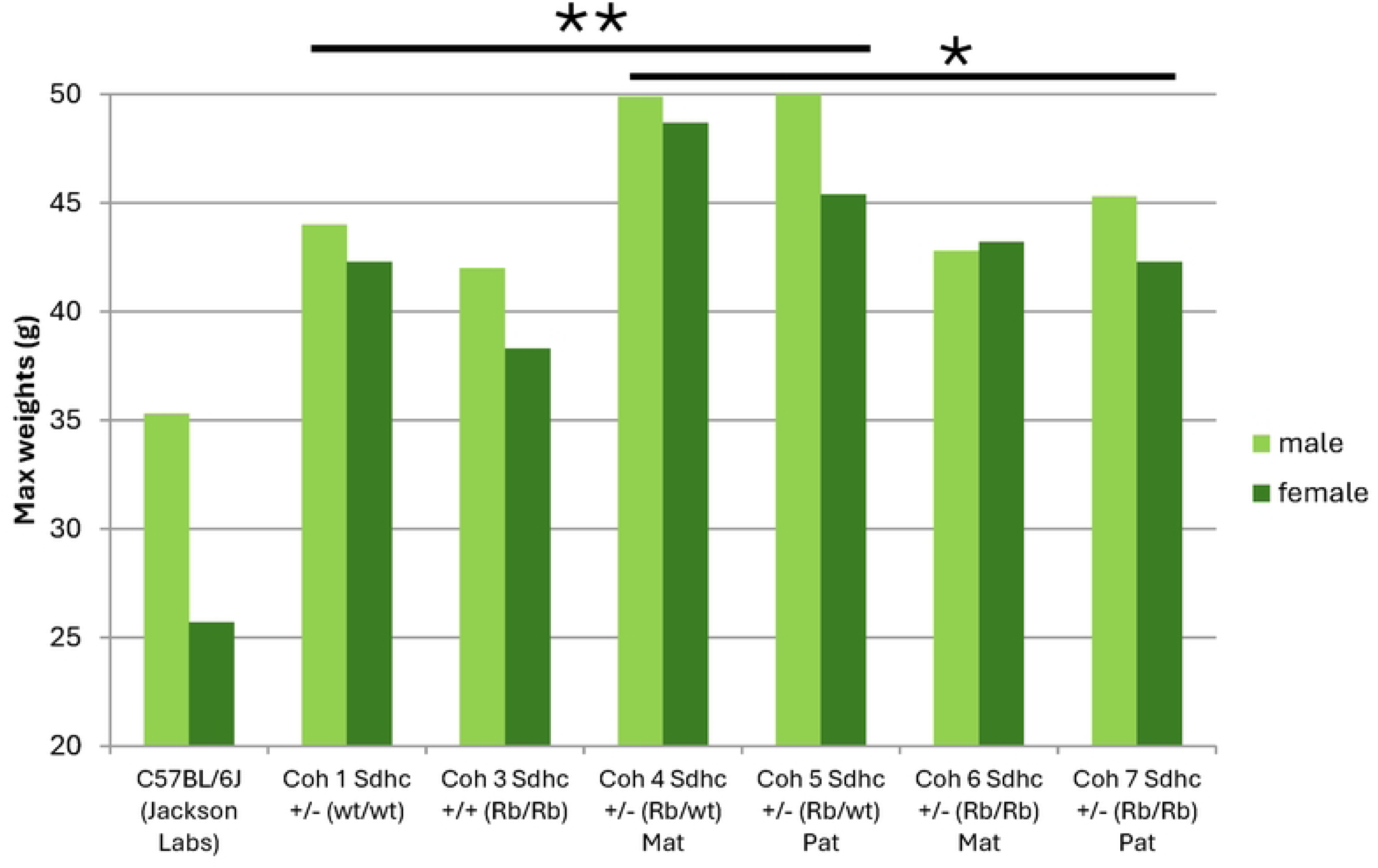
Maximum recorded weight of the various cohorts. Weight of wildtype C57BL/6J (data Jackson Laboratory - left) at 25 weeks included for reference. Mean cohort age was 85 weeks. Cohorts 4 and 5, carrying an Sdhc heterozygous KO and a single Robertsonian chromosome, showed the highest weights (All Coh 4-5 versus Coh 6-7, two-tailed unpaired T test *p=0.005, corrected; All Coh 1-3 versus All Coh 4-5, **p>0.0001). In cohorts 4&5 versus 6&7, males accounted for most of the excess weight (males p=0.01, corrected; females p=NS). Although lower than cohorts 4&5, cohorts 6&7 carrying two Robertsonian chromosomes were significantly heavier compared with cohorts 1-3 (p=0.03, corrected). To avoid confounding effects of pathology on body weight, weights were analyzed as maximum lifetime weight achieved. P values corrected for multiple testing.

### Mouse body weight

The loss of Sdhc is associated with weight gain in mice, as previously reported by the IMPC, which found heterozygous (Sdhc+/-) weights of 43g and 33g for males and females, respectively (n=15). In our cohort Sdhc +/-mice showed similar weights of 44g and 41g for males and females, respectively (n=17). Interestingly, mice in the experimental cohorts 4 and 5, carrying an Sdhc heterozygous KO and a single Rb chromosome, showed the highest weights compared with both the control cohorts (All Coh 1-3 versus All Coh 4&5, **p>0.0001) and the Rb/Rb experimental cohorts (All Coh 4&5 versus Coh 6&7, p=0.005). Nevertheless, cohorts 6&7 were also significantly heavier compared with cohorts 1-3 (p=0.03). In cohorts 4&5 versus 6&7, males accounted for most of the excess weight (males p=0.01; females p=NS). In the experimental cohorts, carrying a single Rb chromosome together with heterozygosity for Sdhc appeared to confer excess body weight in addition to the contribution of Sdhc alone.

### Pathology

In addition to the adrenal gland, the abdominal aortal area and carotid body, the primary sites of paraganglioma-pheochromocytoma in humans, we also assessed general pathology in the various cohorts (Figure 4 or Table 1). A low level of pathology in various organs is common in aged mice. Pathology across all cohorts was broadly similar, with the exception of liver, immune and thyroid abnormalities. Liver pathologies appeared lowest in those cohorts carrying two copies of a Rb chromosome (3, 6 & 7).

**Figure 4.**
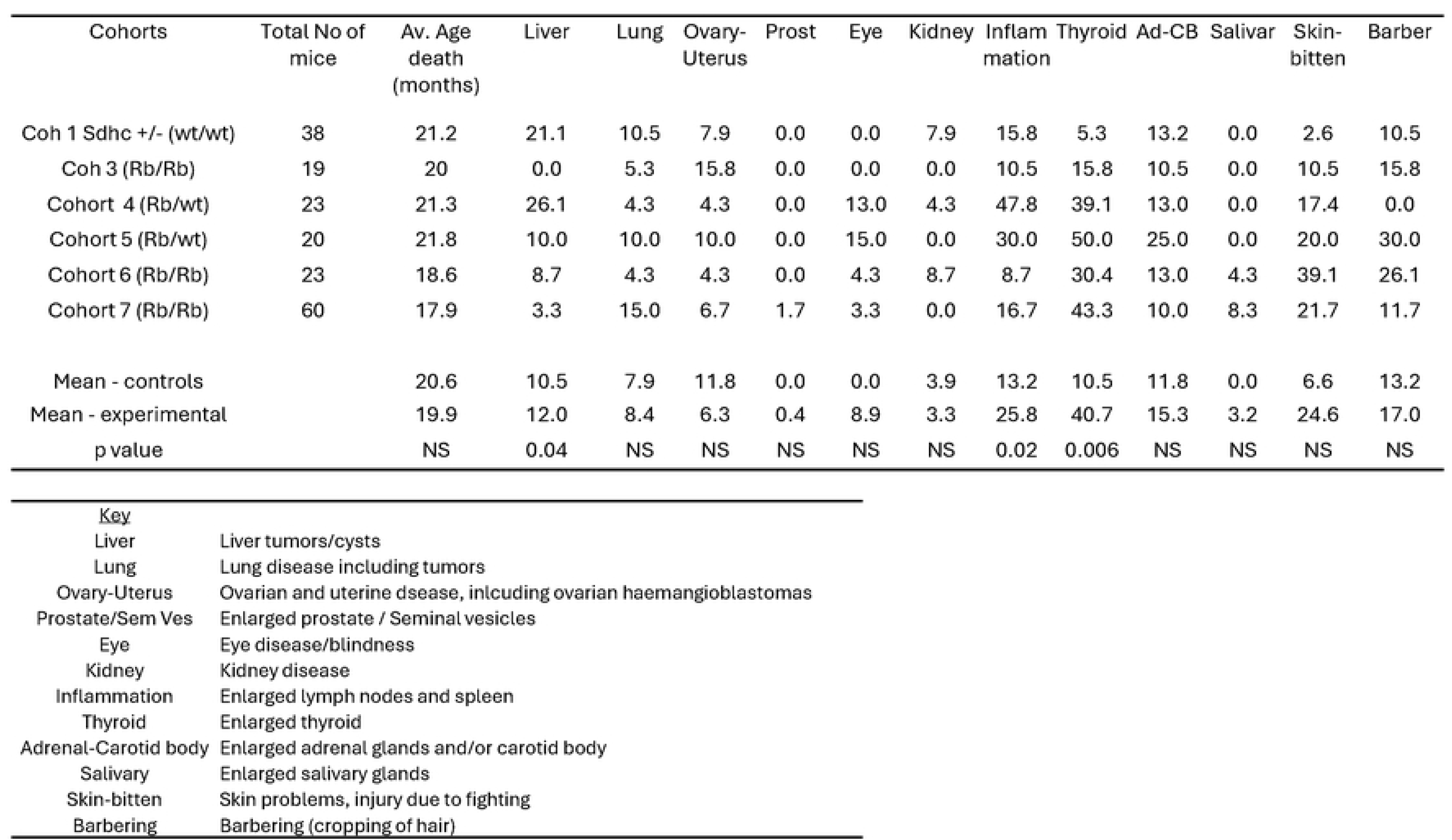
Overview of gross phenotypes and frequencies found in Sdhc +/-Rbl:7 cohorts. Columns indicate organs visually assessed by investigators for signs of pathology. Key provides more detail on pathology observed. Liver tumours/cysts (not further investigated) were marginally more common in cohorts 4 & 5. Inflammation refers to noticeable enlargement of any lymph node and/or spleen, which was significantly more frequent in cohorts 4 & 5 harbouring one copy of the Rbl:7 Robertsonian chromosome. The main finding, beside an incidental bilateral pheochromocytoma, was conspicuous pathology of the thyroid, which was highly significant in the experimental versus control cohorts. P value for cross cohort significance and corrected for multiple testing.

### Inflammation

A second difference between cohorts was the regular occurrence of gross, systemic inflammation in mice carrying a single copy of a Rb chromosome (Figure 4 or Table 1). This was apparent due to grossly enlarged lymph nodes and/or spleen (Figure 5). While this might seem related to skin injuries, self-inflicted or due to fighting, which were broadly more common in experimental versus control cohorts, inflammation did not correlate with either skin injury or with barbering (without skin injury).

**Figure 5.**
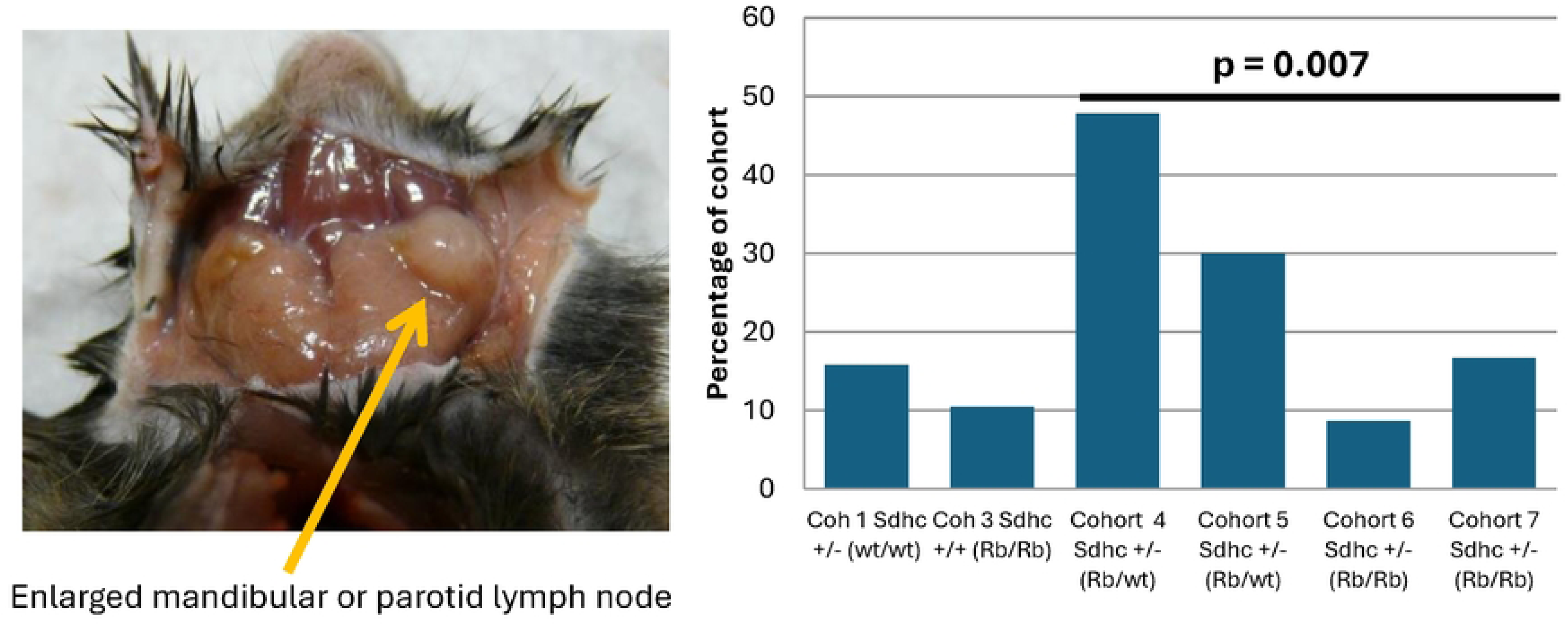
Immune abnormalities were significantly more common in Sdhc +/-Rbl:7 cohorts carrying only one Robertsonian chromosome compared to cohorts with two Robertsonian chromosomes. Immune abnormalities in this cohort did not correlate with fighting or other external injury or with any other pathology. Cross-cohort Chi Sq (corrected) significance p=0.02. Cohorts 4&5 versus cohorts 6&7, 2x2 contingency p=0.007 (corrected). Corrected= correction for multiple testing.

### Thyroid abnormalities

Gross pathology of the thyroid is extremely rare in the C57BL/6N mouse line. The experimental Sdhc+/-Rb1:7 cohorts developed high rates of thyroid disease (Figure 4 or Table 1, Figure 6), with mice exhibiting hyperplasia and papillary thyroid carcinoma-like tumours. Up to 50% of mice in the Rb experimental cohorts exhibited enlarged thyroid glands, which emerged from under the sternothyroid muscle upon necropsy. The low level of abnormalities in the Sdhc+/-cohort and the higher rate in the Rb/Rb (Sdhc +/+) cohort suggest a possible synergistic effect of one or more Rb chromosomes with the presence of heterozygous Sdhc. Histochemical analysis of the thyroid of affected mice showed extensive hyperplasia, cystic areas, and a predominantly papillary morphology in many cases, suggestive of papillary thyroid carcinoma (Figures 7 & 8).

**Figure 6.**
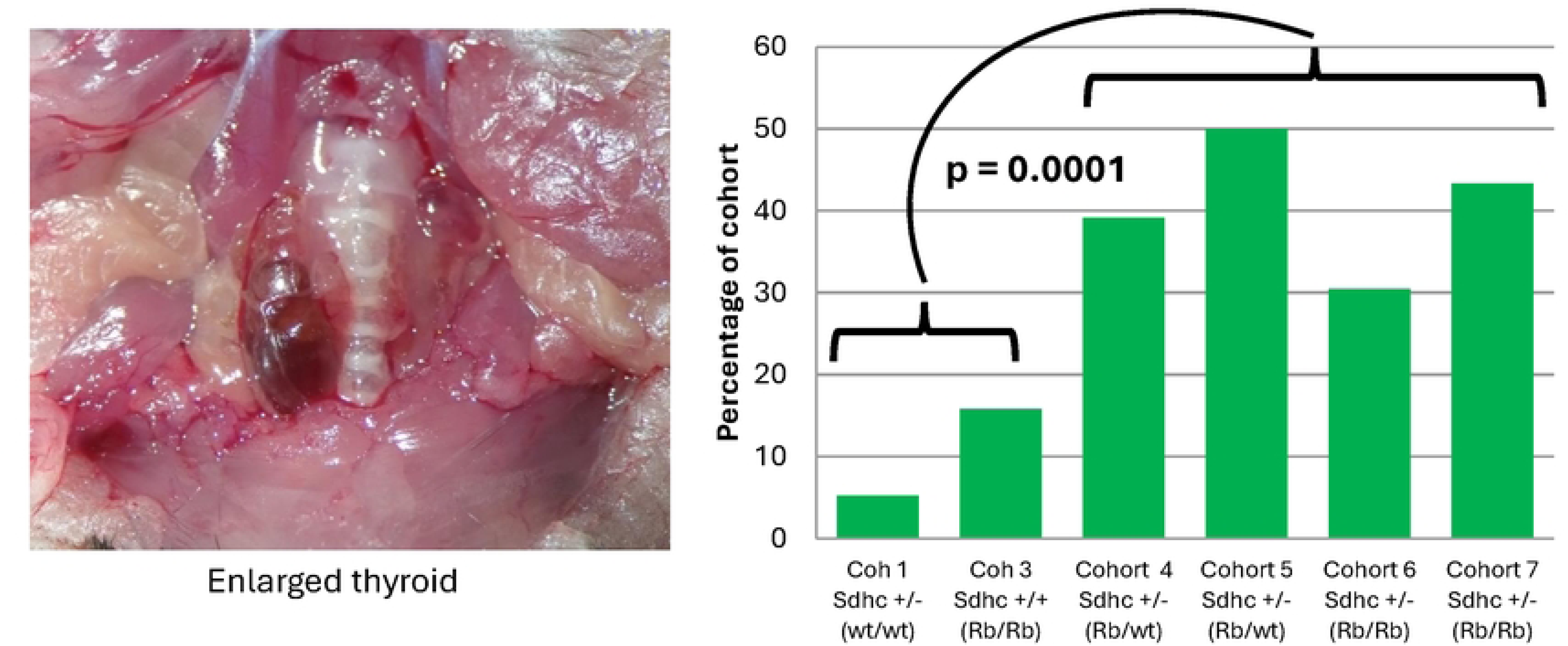
Thyroid abnormalities were common in Sdhc +/-Rbl:7 cohorts. Mouse 9104481 (2.5X objective) showing an enlarged thyroid, visible next to the trachea. In general, gross pathology of the thyroid was rare in the C57BL6 mouse background, as illustrated by the Sdhc+/-cohort (5%, left panel). Cross-cohort Chi Sq significance p=0.006 (corrected). Cohorts 1-3 versus 4-7, p *=* 0.0001 (corrected). Cohorts 4&5 versus 6&7, p=NS. Cohorts 4&6 (maternally transmitted mutation) versus 5&7 (paternally transmitted mutation), p=NS. Corrected= correction for multiple testing.

**Figure 7.**
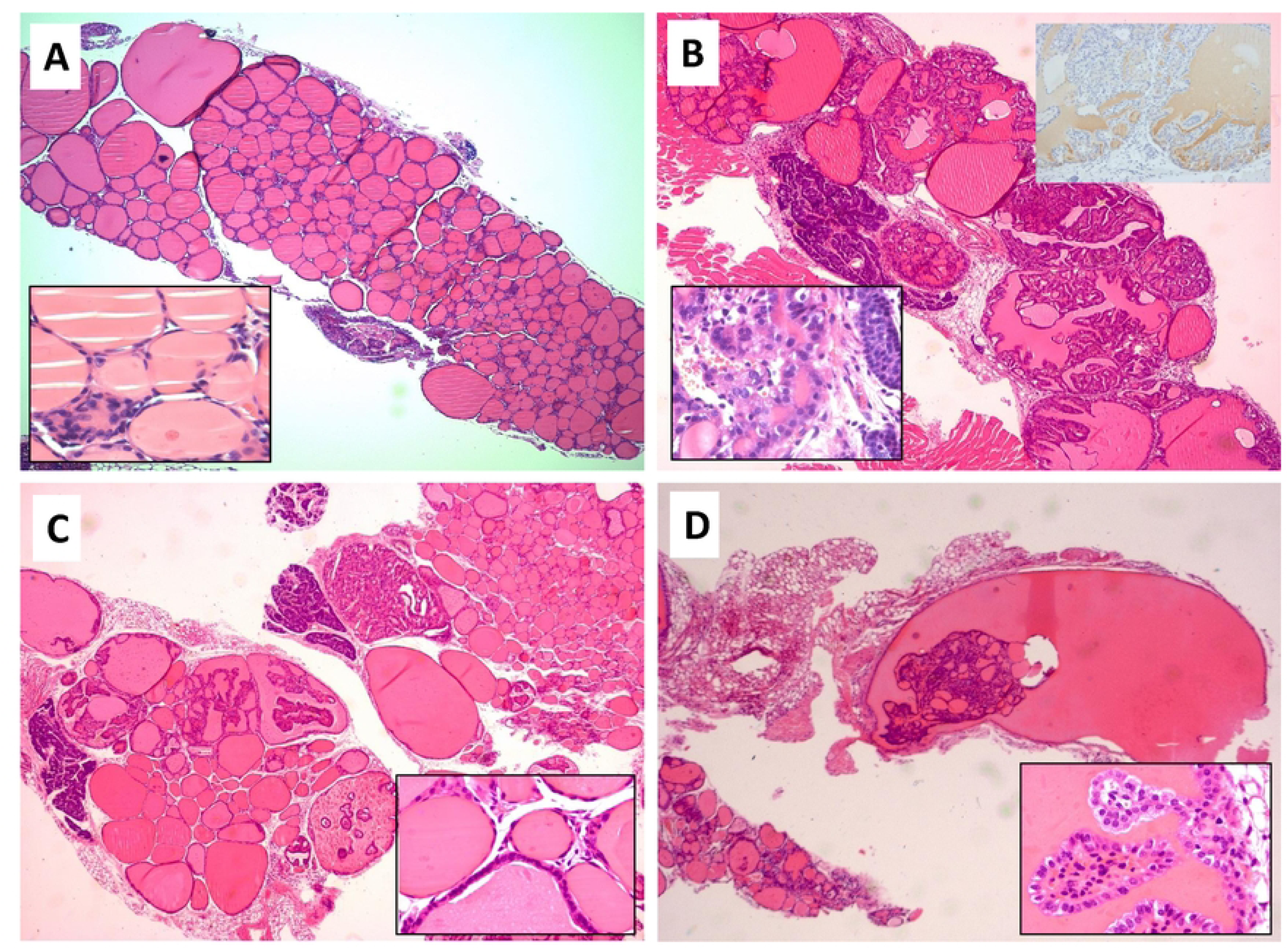
Typical thyroid abnormalities found in cohorts 4 to 7 mice. **(A)** Hematoxylin-eosin histology of thyroid from a healthy wildtype female mouse; 2.Sx objective. Inset 40x objective (resized). **(B)** Thyroid from mouse 9098841 (female, cohort 5 or 7 - metaphase unsuccessful) showing highly abnormal histology, diagnosed as a papillary carcinoma or hyperplasia; 2.Sx objective. Inset lower left H&E 40x objective (resized); inset upper right 40x objective (resized), anti-thyroglobulin IHC. **(C)** Thyroid from 9103313 (female, cohort 7) diagnosed as a papillary carcinoma; 2.Sx objective. Inset 40x objective (resized). **(D)** 9104481 (female, cohort 7) diagnosed as a papillary carcinoma; 2.Sx objective. Inset 40x objective (resized).

**Figure 8.**
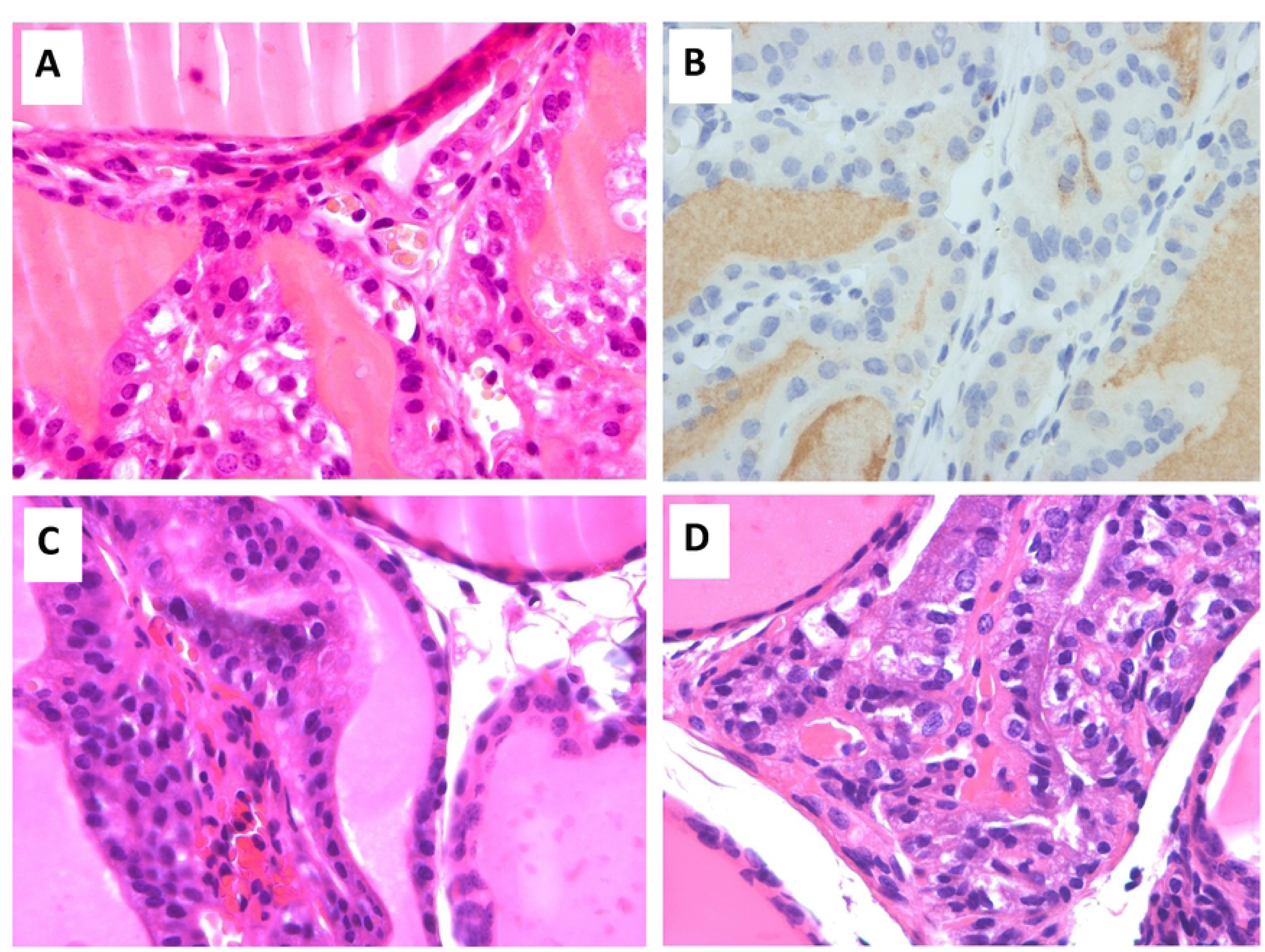
Detailed view of thyroid abnormalities found in cohorts 4 to 7 mice (40x objective). (A) Thyroid from mouse 9098841 (female, cohort 5 or 7 - metaphase unsuccessful) showing highly abnormal histology, including enlarged nuclei and diagnosed as a papillary carcinoma or hyperplasia. (B) Thyroid from mouse 9098841stained with anti-thyroglobulin. (C) Thyroid from 9101254 (female, cohort 5) diagnosed as possible C cell proliferation. (D) Thyroid from 9103110 (female, cohort 5) diagnosed as cystic papillary carcinoma.

### Pheochromocytoma

An interesting incidental finding was bilateral pheochromocytoma in a single mouse (9100515 Sdhc +/-female, cohort 5 or 7 – metaphase not completed). The left-sided tumour showed dramatic enlargement of the adrenal medulla (Figures 9a&b), consistent with a diagnosis of pheochromocytoma (643mg). The right-sided adrenal was also grossly enlarged (Figure 9b) at 18mg, which is 5x normal mean weight. The mean size of the adrenal gland in wildtype C57BL6 is 3.6mg (range 1mg to 5.5mg) (Figure 9c). Immunohistochemical (IHC) staining with anti-tyrosine hydroxylase, a marker protein found in the adrenal medulla and paraganglioma/pheochromocytoma, showed histology typical of pheochromocytoma (Figure 9d). To assess a possible role for loss of Sdhc in the genesis of these tumours, we carried out anti-SDHB IHC, which is usually negative in SDH-associated tumours due to loss of the SDH protein complex. Here, however, staining was positive, suggesting no role for Sdhc loss (Figure 9e). Another characteristic of SDH-deficient tumours is loss of 5-hydroxymethylcytosine (5hmC), as seen in a human SDHB-deficient tumour (Figure 9f). Pheochromocytoma cells from mouse 9100515 showed anti-5hmC staining (Figure 9g) comparable to a wildtype mouse (Figure 9h), suggesting retention of 5hmC and again no causative link to loss of Sdhc. Furthermore, PCR of tumour DNA showed retention of wildtype Sdhc (data not shown), further confirming the lack of a role for SDH loss in genesis of these tumours. Pheochromocytoma cells were cultured but failed to proliferate (Figure 9i).

**Figure 9.**
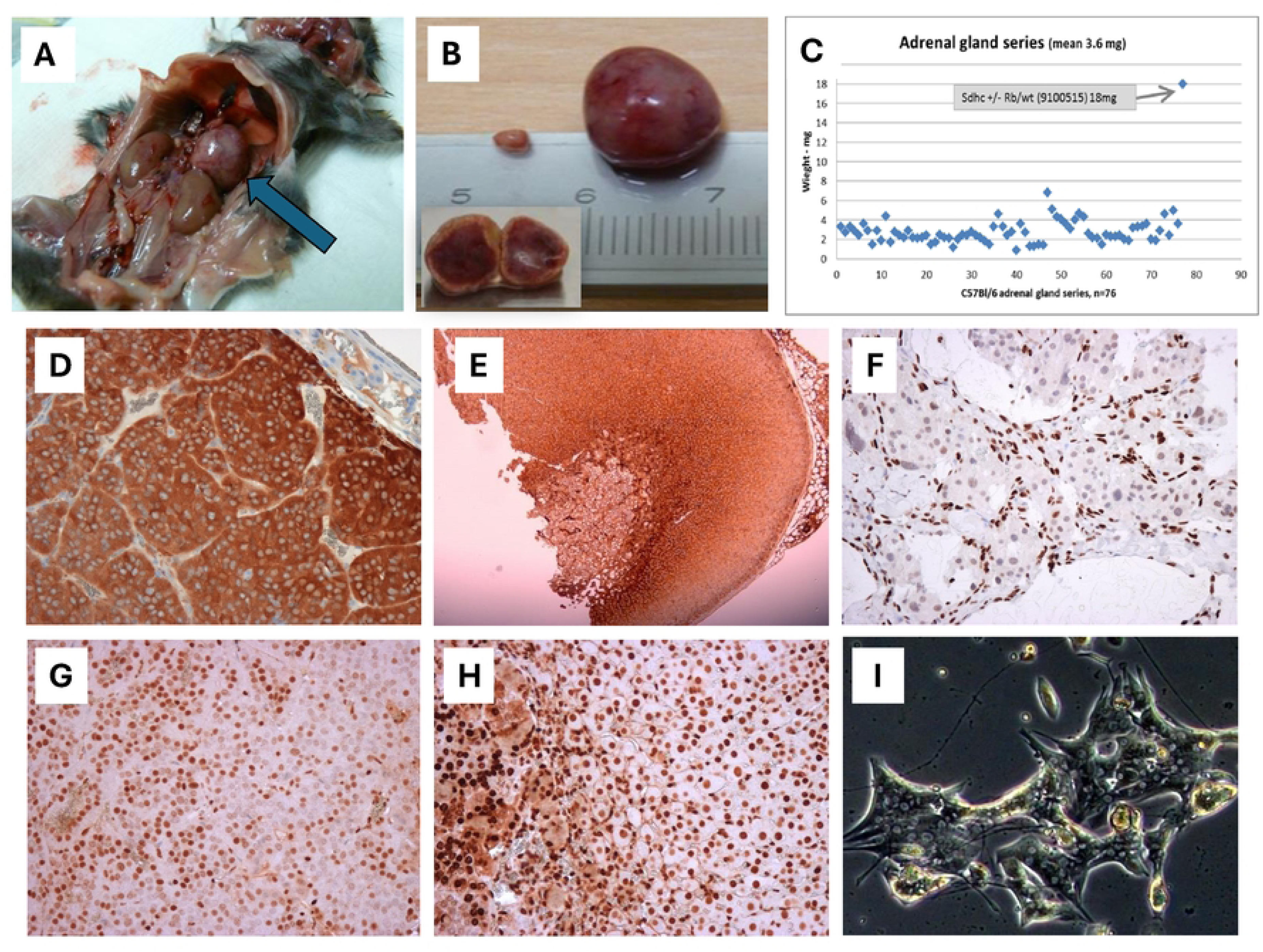
(A) Mouse 9100515 Sdhc +/-Rb/? (female, cohort 5 or 7 - metaphase unsuccessful) shown with *in situ* left-sided pheochromocytoma. The enlarged right­ sided adrenal is also visible. **(B)** The right-sided adrenal weighted 18mg and the left­ sided 643mg. Inset shows cross-section of the larger adrenal. **(C)** A series of 76 adrenal glands from mice with a C57BL/6 background. At 18mg the right-sided adrenal was a major outlier. **(D)** Anti-tyrosine hydroxylase immunohistochemical staining, a marker protein found in the adrenal medulla and pheochromocytomas, shows typical histology in the 9100515 pheochromocytoma. **(E)** Anti-SDHB staining is generally negative in SDH-associated tumours due to loss of the SDH complex, but is positive here, suggesting a tumour unrelated to Sdhc loss. **(F)** Another characteristic of SDH-deficient tumors is loss of 5-hydroxymethylcytosine (5hmC) as seen here in a human SDHB­ deficient tumour. **(G)** Anti-5hmC staining of 9100515 pheochromocytoma cells compared with a wildtype mouse **(H)** showed retention of 5hmC. PCR of tumour DNA also showed retention of wildtype Sdhc. (I) 9100515 pheochromocytoma cells in culture showed unusual morphology but failed to proliferate.

## Discussion

This study had two primary purposes: 1) to test the Hensen model of paternal inheritance of SDH-related tumour risk, and 2) to develop a paraganglioma mouse model to study the genesis of paraganglioma. While not confirmed in this study, neither is the Hensen model refuted, and it remains the most compelling explanation of the available evidence on Sdh paternal inheritance [13]. Numerous groups over the past two decades have sought to create an Sdh-related paraganglioma mouse model [14]. None have been successful to date. A third goal of this study was to explore the feasibility of investigating genetic modifiers using non-standard chromosomal configurations in the mouse. In this case, the objective was partly successful, in that a combination of a heterozygous genetic knockout and a Rb chromosome produced a stronger phenotype compared to the Rb/Rb or Sdhc +/-cohorts alone (50% vs. 15% or 5%).

### A human semi-homologous chromosome in mice

Rb translocations in mice have been studied extensively but mainly in relation to processes that result in chromosomal races, reproductive isolation and speciation. Phenotypically, Rb chromosomes are sometimes associated with reduced fertility [11], and have been associated with differences in growth of the mouse mandible [15], but few other phenotypic traits of wild or laboratory-housed Rb mice have been reported. Karyotype analyses of human *SDHD* and *SDHAF2* tumours invariably show loss of the maternal chromosome [16-19] and the few cases of maternal inheritance described show mitotic recombination and subsequent loss of the short arm of maternal chromosome 11 [20, 21], underlining the essential role of this chromosomal region in tumourigenesis. The strategy adopted in the present study was chosen because the modifier required for tumour initiation in human carriers of *SDHD* and *SDHAF2* has not been precisely defined [22]. It is also possible that more than one gene is involved, at least one of which is likely imprinted on the paternal chromosome and expressed from the maternal chromosome. While our strategy brought together homologous chromosomal regions essential for SDH-initiated tumourigenesis, weaknesses of this approach are that homologous regions are absent and in the event of an LOH event non-homologous regions will be lost, potentially causing or inhibiting emergence of phenotypes.

### Inflammation

A novel observation in this study was the high incidence of gross inflammation in mice carrying only one copy of an Rb 1:7 chromosome. These mice were characterized by grossly enlarged lymph nodes in a variety of locations, and in some cases the spleen was enlarged. Both the maternal and paternal cohorts 4 and 5, each of which carried one metacentric 1:7 chromosome (together with telocentric chromosomes 1 and 7), showed a significantly higher frequency of gross inflammation upon necropsy compared to Sdhc+/-mice carrying two Rb chromosomes or the control cohorts. The absence of similar findings in the Rb/Rb cohorts 6 and 7 suggests that Sdhc did not play a role in this phenotype. Heterozygosity for a metacentric chromosome has previously been reported to influence some phenotypes [23] but immune anomalies have not been reported to our knowledge.

### Thyroid

With an incidence of 1%, thyroid carcinoma is the most common endocrine cancer, and the subtype papillary thyroid carcinoma (PTC) is the most frequent variety, accounting for around 80% of all thyroid cancers. PTCs are a minor diagnostic criterium of Cowden Syndrome and have been reported in isolation in association with PPGL [24, 25], in association with *SDHB* [26], *SDHC* [27-29], *SDHD* [30-33] and *SDHAF2* [34] variants. The lower level of thyroid abnormalities in Sdhc +/-mice compared to Rb/Rb Sdhc +/+ suggest that the Robertsonian chromosome was the primary contributor to the phenotype, but the high levels in the combined Rb/Sdhc+/-cohort indicate potent synergism.

As mentioned, missense variants in succinate dehydrogenase (SDH) complex genes have been previously linked to human thyroid cancer. To explore this subject further, Ashtekar and colleagues generated mice lacking Sdhd in the thyroid. These mice developed enlarged thyroid glands with follicle hypercellularity and increased proliferation. In addition, knockdown of *SDHD* in human thyroid cell lines caused enhanced migratory capability and stem-like features, which were also observed in the mouse tumours. These characteristics could be reversed by α-ketoglutarate, suggesting dedifferentiation occurs due to an imbalance in TCA metabolites [35]. In a study focussed on Cowden Syndrome patients, Ni and colleagues found that both papillary and follicular thyroid tumours showed consistent loss of SDHC/D gene expression, which was associated with earlier disease onset and higher pathological-TNM stage [29].

### Pheochromocytoma

Another unusual finding of the study was the occurrence of a pheochromocytoma in a mouse from the experimental cohort 5/7 (metaphase unsuccessful), the experimental cohorts designed to directly mimic the Hensen model. While many rodent models develop adrenal pheochromocytomas, including Nf1 knockouts [36], c-Mos transgenics [37], RET Met918 transgenics [38], Cdkn1b-mutated Sprague–Dawley rats [39], Rb1/Trp53 dual knockouts [40], ceramide synthase 2 knockout mice [41], ErbB2 transgenics [42], connexin 32 knockouts [43], PTEN knockouts [44] and B-Raf transgenics [45], spontaneous pheochromocytoma in the C57BL/6 mouse background has not been reported to our knowledge. The co-occurrence of a pheochromocytoma with a smaller tumour/hyperplasia therefore suggests a germline or very early genetic event. However, loss of Sdhc did not appear to play a role.

## Conclusion

As both the present mouse model, as well as our previous Sdhd/H19 KO, did not develop paragangliomas, the Hensen model remains hypothetical. A conclusive test of the Hensen model will likely require an animal model that regularly develops paraganglioma. The striking phenotypes reported here highlight the lack of Robertsonian studies to date that considered phenotype or pathology. We further conclude that chromosomal constitution can dramatically impact phenotype. The approach chosen here, if more widely applied, may aid in the dissection of other diseases in which modifiers are suspected.

## Contributor Role

Conceptualization: Jean-Pierre Bayley

Data Curation: Jean-Pierre Bayley, Heggert Rebel

Formal Analysis: Jean-Pierre Bayley, Heggert Rebel

Funding Acquisition: Jean-Pierre Bayley

Investigation: Jean-Pierre Bayley, Heggert Rebel

Methodology: Jean-Pierre Bayley, Peter Devilee

Project Administration: Jean-Pierre Bayley, Peter Devilee

Resources: All authors

Supervision: Jean-Pierre Bayley, Peter Devilee

Validation: Jean-Pierre Bayley, Peter Devilee

Visualization: Jean-Pierre Bayley, Heggert Rebel

Writing – Original Draft Preparation: Jean-Pierre Bayley

Writing – Review & Editing: All authors

## Funding

This work was supported by the Dutch Cancer Foundation (KWF UL 2011-5025) and The Paradifference Foundation. No funding source played any role in the design, analysis, writing of the manuscript or the decision to submit for publication.

## Competing interests

The authors have declared that no competing interests exist.

## Acknowledgements

We thank Professor Judith Bovee for assistance with thyroglobulin staining, and Professor Hans Morreau and Dr. Daniela Salvatori for assessment of thyroid pathology.

## Data Availability Statement

For full data access please apply to corresponding author.

## Statement of Ethics Approval

This study was approved by the Animal Experimentation Committee of Leiden University Medical Center (DEC nr12069).

## Notes

### Competing Interest Statement

The authors have declared no competing interest.

